# MHC Molecules on B Cell Microvilli Are Spatially Associated with IL-15Rα

**DOI:** 10.64898/2026.09.24.747988

**Authors:** István Rebenku, Noémi Bilakovics, Levente Szolyka, Lukas Velas, Tamás Garda, Gerhard J. Schütz, György Vámosi

## Abstract

Interleukin-15 (IL-15) trans-presentation (TP) by B cells is an important mechanism of T-cell activation; however, the spatial organisation of interleukin-15 receptor α (IL-15Rα) relative to major histocompatibility complex (MHC) molecules on B-cell microvilli remains poorly understood. As microvilli protrude from the B-cell surface and may serve as sites of initial B-cell–T-cell contact, the distribution of IL-15Rα and MHC molecules within these structures may contribute to the earliest stages of T-cell recognition and activation. Here, we investigated the spatial association and molecular proximity of IL-15Rα with MHC class I and class II molecules on B-cell microvilli before immunological synapse formation, using confocal microscopy, stimulated emission depletion (STED) microscopy, stochastic optical reconstruction microscopy (STORM), and fluorescence lifetime imaging microscopy–based Förster resonance energy transfer (FLIM-FRET). Both MHC class I and MHC class II molecules showed a significant spatial association with IL-15Rα; however, the extent of colocalisation decreased as the spatial resolution increased. STED microscopy revealed significant colocalisation between IL-15Rα and MHC class I, whereas STORM did not detect this association. In contrast, IL-15Rα and MHC class II remained significantly colocalised at both resolutions. FLIM–FRET further demonstrated molecular proximity between IL-15Rα and both MHC class I and MHC class II molecules, with higher FRET efficiency observed for MHC class II. Collectively, these findings indicate that IL-15Rα is spatially organised in proximity to both MHC class I and class II molecules on B-cell microvilli before immunological synapse formation. This arrangement at potential sites of initial B cell–T cell contact may facilitate the coordination of IL-15 trans-presentation and antigen presentation during the earliest stages of B cell–T cell interactions.

## Introduction

Microvilli are actin-rich, finger-like protrusions found on T lymphocytes, B lymphocytes and dendritic cells [1–3]. These dynamic structures are enriched in antigen-presenting, antigen-recognising, and co-stimulatory molecules, including major histocompatibility complex class II (MHC class II), the T-cell receptor (TCR), cluster of differentiation 4 (CD4), cluster of differentiation 2 (CD2), intercellular adhesion molecule 1 (ICAM-1), C–C chemokine receptor type 7 (CCR7), and lymphocyte function-associated antigen 1 (LFA-1) [1, 4–8]. The clustering of signalling molecules increases signalling efficiency [9]. Microvilli can penetrate the glycocalyx, thereby reducing the intercellular distance between cells [10] and establishing initial sites of cell–cell contact. This process is essential for immunological synapse formation [6].

Interleukin-15 (IL-15) is a pleiotropic cytokine that plays a crucial role in inflammation by modulating the responses of both innate and adaptive immune cells. IL-15 promotes the proliferation, activation and survival of natural killer (NK) cells, natural killer T (NKT) cells, CD8+ effector-memory T cells and central-memory T cells. The IL-15 receptor (IL-15R) is a heterotrimeric membrane receptor consisting of three subunits: the ligand-specific IL-15Rα subunit, the interleukin-2/15 receptor β (IL-2/15Rβ) subunit shared with interleukin-2 (IL-2) and the common γ_c_ chain.

IL-15 is linked to the development of autoimmune diseases [11]. In autoimmune hepatitis (AIH), B-cell depletion improves liver function and reduces the number of CD8+ T cells in the liver. RNA sequencing of these aberrant B cells has shown that IL-15 overexpression is a central driver of their pathogenic activity. In addition, IL-15 expression in B cells is promoted by CD40L+ CD8+ T cells, suggesting mutual interactions between these cell populations [12]. Another study demonstrated that, in primary biliary cholangitis, IL-15Rα+ B cells are enriched around bile ducts in the liver and secrete chemokine C-C motif ligand 3 (CCL3). CCL3 recruits C-C chemokine receptor type 5-positive (CCR5+) CD4+ tissue-resident memory T cells and enhances their effector function through IL-15 trans-presentation. Furthermore, inhibition of IL-15 signalling reduces liver injury in mouse models [13].

Trans-presentation (TP) is a unique mode of cytokine signalling and the principal mechanism of IL-15 action. During this process, an antigen-presenting cell (APC) displays IL-15 complexed with IL-15Rα and presents the cytokine to the IL-2/15Rβ–γ_c_ complex expressed on recipient T or NK cells [11, 14]. Previously, our research group used biophysical methods to demonstrate the intercellular assembly of IL-15R subunits during IL-15 TP at the immunological synapse (IS) formed between Raji B cells and Jurkat CD4+ T cells. We also investigated the relationship between TP and antigen presentation (AP). We found that MHC class II and IL-15Rα are molecular neighbours in B cells and translocate together to the IS in the presence of ligands, including IL-15 and/or superantigen. At the IS, they form intercellular interactions with the TCR/CD3 complex and the IL-2/15Rβ complex, respectively. Formation of the antigen-presenting complex did not affect IL-15 TP-induced STAT5 phosphorylation, suggesting that IL-15 TP can occur autonomously or together with AP [15]. On the other hand, T-cell activation was significantly enhanced *in vivo* when mice were treated with soluble IL-15 [16]. We also found that IL-15Rα colocalised with MHC class I or class II molecules in lipid rafts of T-lymphoma cells [17, 18]. Although microvilli have been extensively studied on T cells [6], their structure and function on B cells remains unclear. We therefore aimed to quantitatively assess the interactions of IL-15Rα with MHC class I and class II molecules on isolated B-cell microvilli before IS formation. To this end, we used microscopy-based approaches, including confocal microscopy, stochastic optical reconstruction microscopy (STORM), stimulated emission depletion (STED) microscopy and fluorescence lifetime imaging microscopy–based Förster resonance energy transfer (FLIM-FRET). These methods enabled the morphological identification of microvilli, allowing compartment-specific quantification of pixel-based colocalisation and measurement of molecular proximity by FRET.

## Results

### Pixel-based colocalisation measurements by confocal and STED microscopy

To assess whether IL-15Rα colocalises with MHC class I or class II molecules in the microvilli of Raji cells, we used confocal and STED microscopy followed by correlation analysis. HaloTag–IL-15Rα was labelled with HaloTag STAR RED ligand in Raji cells expressing HaloTag–IL-15Rα, whereas MHC class I and class II molecules were targeted using STAR 580-conjugated W6/32 and L243 Fab fragments, respectively. As a positive control, the heavy chain and light chain (β2m) of MHC class I were labelled with STAR 580- and STAR RED-conjugated W6/32 and L368 Fab fragments, respectively. As a negative control, transferrin receptors, which are enriched in coated pits [19], and MHC class I molecules, which localise to lipid rafts [20, 17], were labelled with STAR 580-conjugated MEM75 Fab fragments and STAR RED-conjugated W6/32 monoclonal antibodies (mAbs), respectively.

Intensity-based colocalisation was quantified with Intensity-based colocalisation within microvilli was quantified using Pearson’s correlation coefficient. Representative images and the results of the correlation analysis are shown in Fig. 1. Cell-by-cell Pearson’s correlation coefficient values (*r* ± SD) are presented in Fig. 1B and Fig. 1D.

**Fig. 1.**
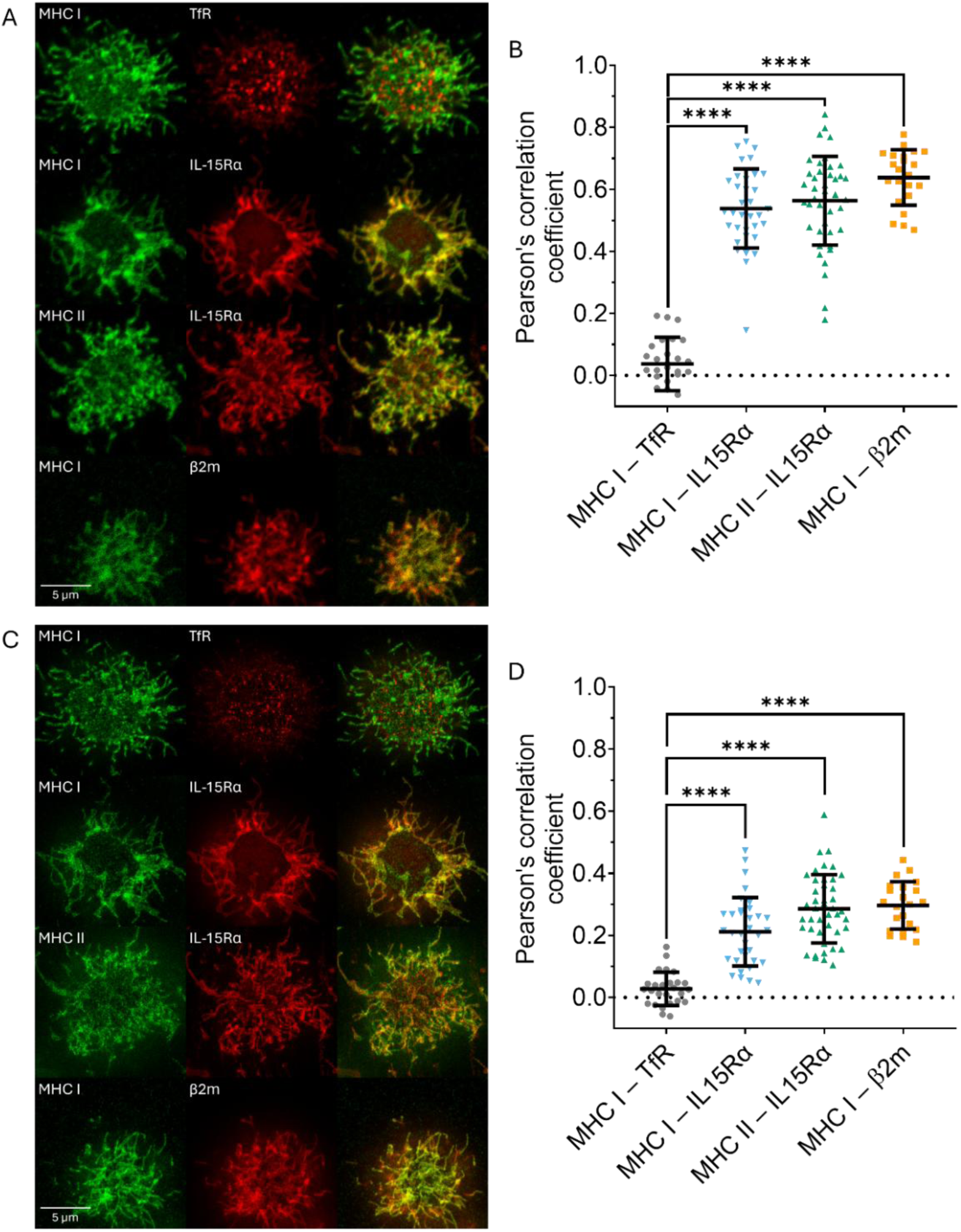
Pixel-based colocalisation measured by confocal and STED microscopy. Optical sections of selected Raji-HaloTag–IL-15Rα cells imaged by confocal (**A**) and STED (**C**) microscopy. The left column shows the farred channel (STAR RED, appearing in red false colour), the middle column shows the red channel (STAR 580, appearing in green false colour), and the right column shows their overlay. Scale bar: 5 µm. (**B**, **D**) Pearson’s correlation coefficients (*r*) between the subcellular intensity distributions of the indicated protein pairs, derived from confocal or STED images. Each dot represents the *r* value of a single cell. Horizontal lines indicate the mean ± SD of the samples. The significance of differences was assessed using a heteroscedastic one-way ANOVA with Welch’s correction; **** p < 0.0001. N = 22-43 cells from n = 4 biological replicates were measured for each protein pair.

Pearson’s correlation coefficients derived from confocal images were 0.54 ± 0.13 and 0.56 ± 0.14 (mean ± SD) for the IL-15Rα–MHC class I and IL-15Rα–MHC class II pairs, respectively, indicating that these proteins were colocalised at the resolution of confocal microscopy (approximately 200 nm). The negative and positive controls yielded values of 0.04 ± 0.09 and 0.64 ± 0.09, respectively.

STED microscopy yielded lower Pearson’s correlation coefficients for all protein pairs: 0.21 ± 0.11 for IL-15Rα–MHC class I, 0.29 ± 0.11 for IL-15Rα–MHC class II, 0.03 ± 0.05 for the negative control and 0.30 ± 0.08 for the positive control. These lower values are attributable to the higher spatial resolution of STED microscopy (approximately 54 nm) and reflect the fact that two proteins cannot occupy precisely the same spatial position. Taken together, these results indicate that IL-15Rα colocalises with both MHC class I and MHC class II molecules in the microvilli of Raji B cells, even at the resolution afforded by STED microscopy.

### Coordinate-based Colocalisation on STORM images

HaloTag FLUX 640 ligand was used to label Halo-IL-15Rα, whilst MHC class I and class II molecules were labelled with Cy3B-conjugated W6/32 and L243 fragments. As a negative control, MHC class I labelled with Alexa Fluor 647-conjugated (abbreviated as A647) W6/32 Fab, paired with transferrin receptor targeted by A647-conjugated MEM75 mAb, was used. MHC class II labelled with Cy3B-L243 Fab and MHC class I labelled with A647-W6/32 Fab served as a positive control, as MHC class I and class II molecules are partially co-clustered in the plasma membrane of JY B and HUT102 T cells [21].

Representative images of the protein localisations are shown in Fig. 2A, and cell-by-cell coordinate-based colocalisation indices are presented in Fig. 2B. The coordinate-based colocalisation (CBC) index values obtained for the MHC class I–MHC class II pair (0.21 ± 0.04) were significantly higher than those of the negative control, MHC class I–TfR (0.05 ± 0.02). Colocalisation index values for the MHC class I–IL-15Rα pair showed no significant difference from the negative control (0.04 ± 0.05), whereas those for the MHC class II–IL-15Rα pair were significantly higher (0.09 ± 0.05).

**Fig. 2.**
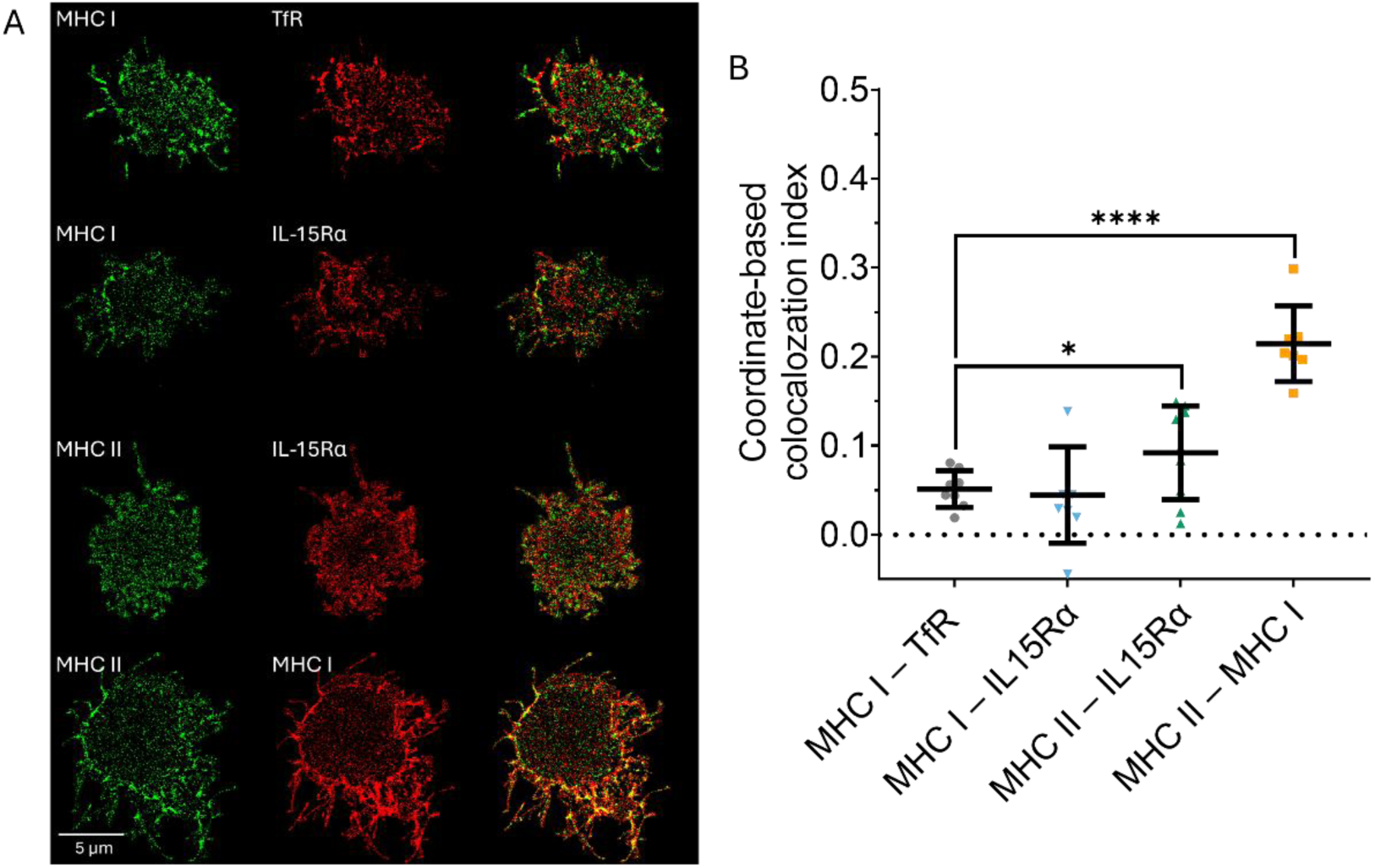
Coordinate-based colocalisation measured via STORM microscopy. **(A)** The left column shows the localisations in the far-red channel (red false colour), the middle column shows the green channel, and the right column shows their overlay. Scale bar: 5 µm. (**B**) Each point represents the coordinate-based colocalisation index of a single cell. Horizontal lines indicate the mean ± SD of the samples. For statistical analysis, Brown–Forsythe and Welch’s heteroscedastic one-way ANOVA were used; * p < 0.05; **** p < 0.0001. N = 7-9 cells were measured.

### FLIM-FRET reveals molecular proximity between IL-and MHC class I and class II molecules in microvilli

Whereas STED and STORM microscopy can directly visualise colocalisation at resolutions of a few tens of nanometres, they cannot detect proximity at the molecular scale. To assess whether IL-15Rα associates with MHC class I or class II molecules in the microvilli of live Raji cells, we performed FLIM-FRET measurements. IL-15Rα on HaloTag–IL-15Rα Raji cells was labelled with the cell-permeable HaloTag TMR ligand, which was used as a FRET donor. A647-conjugated antibodies targeting selected protein partners served as FRET acceptors. Fig.3A shows representative confocal images of selected cells, where the FRET donor (Fig.3A, 1^st^ column), the acceptor (Fig.3A, 2^nd^ column) and their merged view (Fig.3A, 3^rd^ column) can be seen. FRET efficiencies were quantified within microvilli; representative FRET efficiency maps and corresponding histograms are shown in Fig. 3A, 4^th^ and 5^th^ columns, respectively. The fitted fluorescence lifetime decay curves of the selected cells are shown in Fig. S1. Cell-by-cell average (E ± SD, %) values are presented in Fig. 3B.

**Fig. 3.**
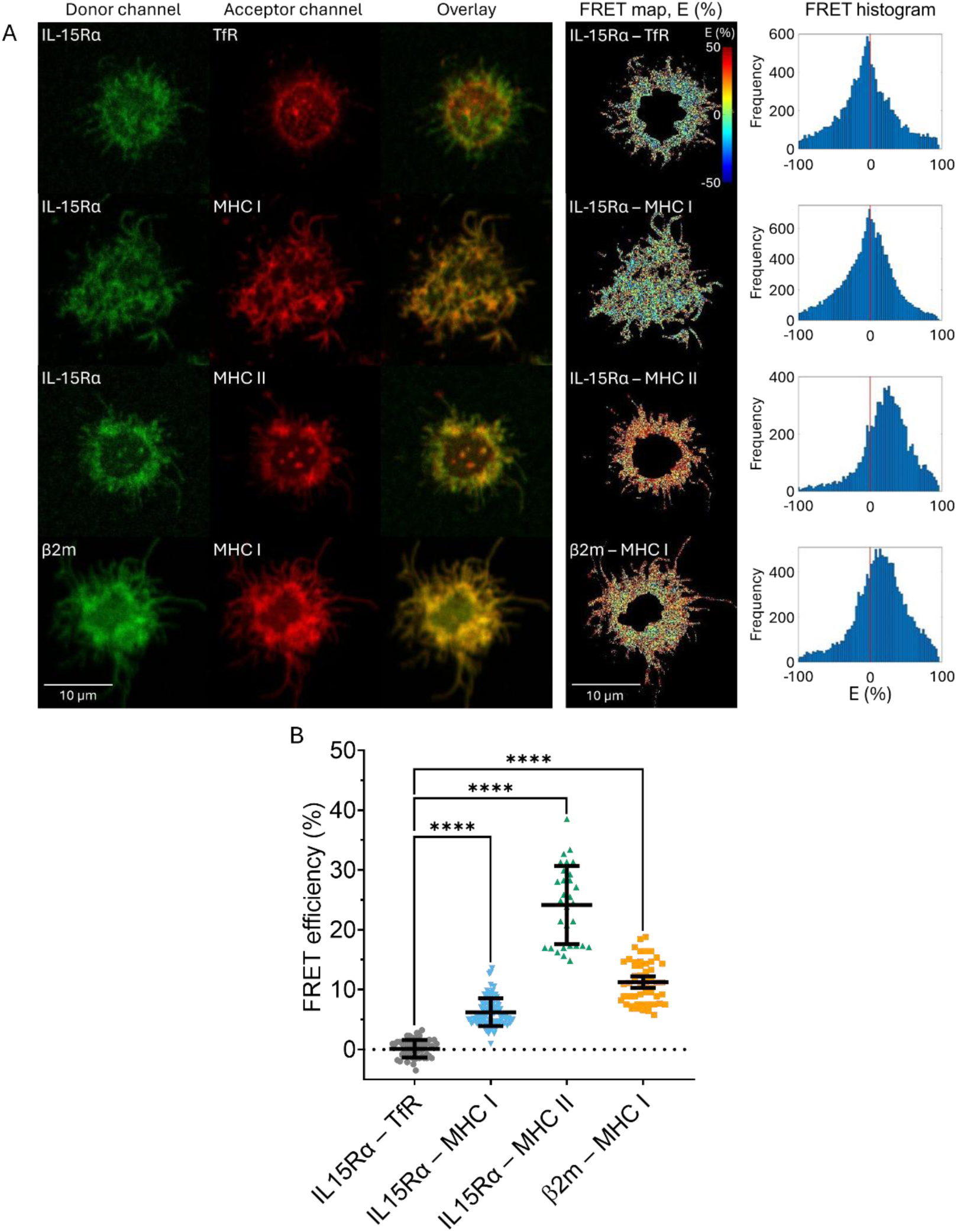
FLIM-FRET analysis of the interaction between IL-15Rα and MHC class I or class II molecules in the microvilli of Raji-HaloTag–IL-15Rα cells. (A) Confocal images of the FRET donor (first column), acceptor (second column), merged channels (third column), FRET-efficiency maps (fourth column) and the corresponding FRET-efficiency histograms (fifth column) of selected Raji-HaloTag–IL-15Rα cells. First row: a cell labelled with HaloTag TMR ligand and A647-MEM75 (anti-transferrin receptor) antibody; second row: a cell labelled with HaloTag TMR ligand and A647-W6/32 (anti-MHC class I) antibody; third row: a cell labelled with HaloTag TMR ligand and A647-L243 (anti-MHC class II) antibody; fourth row: a cell labelled with A546-W6/32 and A647-L368 (anti-β2m) mAbs. Scale bar: 10 µm. (B) Each point represents the mean FRET efficiency of a single cell. Horizontal lines indicate the mean ± SD of the samples (*E* ± SD, %). Group differences were assessed using the Kruskal–Wallis test followed by Dunn’s post hoc test; **** p < 0.0001. N = 30-99 cells from n = 3 biological replicates were measured.

As a negative control, we measured FRET between HaloTag TMR–IL-15Rα and A647-tagged MEM75 antibodies targeting transferrin receptors. The FRET efficiency in these cells was *E* = 0.1 ± 1.4%. In the positive-control sample, the FRET efficiency between the heavy and light chains of MHC class I, labelled with Alexa Fluor 546 (abbreviated as A546)-W6/32 and A647-L368 mAbs, was *E* =11.4 ± 3.4%, consistent with their established molecular association [22]. FRET measurements between TMR-labelled IL-15Rα and MHC class I, labelled with A647-W6/32 mAb, yielded *E* = 6.2 ± 2.3%, which was significantly higher than that of the negative control. The highest FRET efficiency, *E* = 24.1 ± 6.5%, was observed between IL-15Rα and MHC class II. These results indicate that IL-15Rα associates with both MHC class I and class II molecules on the microvilli of Raji B cells. Notably, the substantially higher FRET efficiency observed with MHC class II suggests a preferential association of IL-15Rα with MHC class II in these protrusions.

## Discussion

Antigen presentation and IL-15 trans-presentation represent distinct, though potentially related, modes of communication between antigen-presenting cells and T cells. We previously demonstrated, using a Raji B-cell– Jurkat T-cell model, that these processes may occur either simultaneously or independently. In that study, IL-15Rα and MHC class II were found to be associated within the Raji-cell plasma membrane and to undergo coordinated translocation to the immunological synapse during both antigen presentation and IL-15 trans-presentation. Moreover, IL-15 trans-presentation and MHC class II-mediated antigen presentation were shown to enhance one another’s signalling in T cells [15]. Consistent with the possibility of a common membrane organisation, IL-15Rα, MHC class I and MHC class II molecules have also been reported to associate within supramolecular membrane clusters and lipid-raft domains [17, 18, 20, 21]

Microvilli are actin-supported membrane protrusions that substantially increase the accessible cell-surface area and have emerged as important sites for environmental sensing and the initiation of lymphocyte cell–cell interactions [1–3, 9]. In T cells, TCR, CCR7 and LFA-1 have been reported to be enriched on microvilli, consistent with a role for these structures in the initial stages of T-cell recognition and adhesion [5, 7, 8]. The presence of signalling and adhesion molecules on microvilli raises the possibility that these protrusions constitute specialised membrane platforms that facilitate the earliest molecular encounters between interacting lymphocytes [6, 10]. Whether a corresponding organisation exists on B-cell microvilli, particularly for molecules involved in antigen presentation and cytokine trans-presentation, has remained unexplored.

In the current study, using a combination of multiscale fluorescence microscopy approaches, we investigated the spatial organisation of IL-15Rα and MHC class I and class II molecules on B-cell microvilli before immunological synapse formation. The complementary microscopy approaches used in this study probe different spatial scales, ranging from membrane-level co-distribution assessed by confocal and STED microscopy to nanoscale co-organisation examined by single-molecule localisation microscopy and molecular proximity evaluated by FRET (Fig. 4). Together, these measurements provide an increasingly detailed description of the spatial relationship between IL-15Rα and MHC molecules on the B-cell surface.

**Fig. 4.**
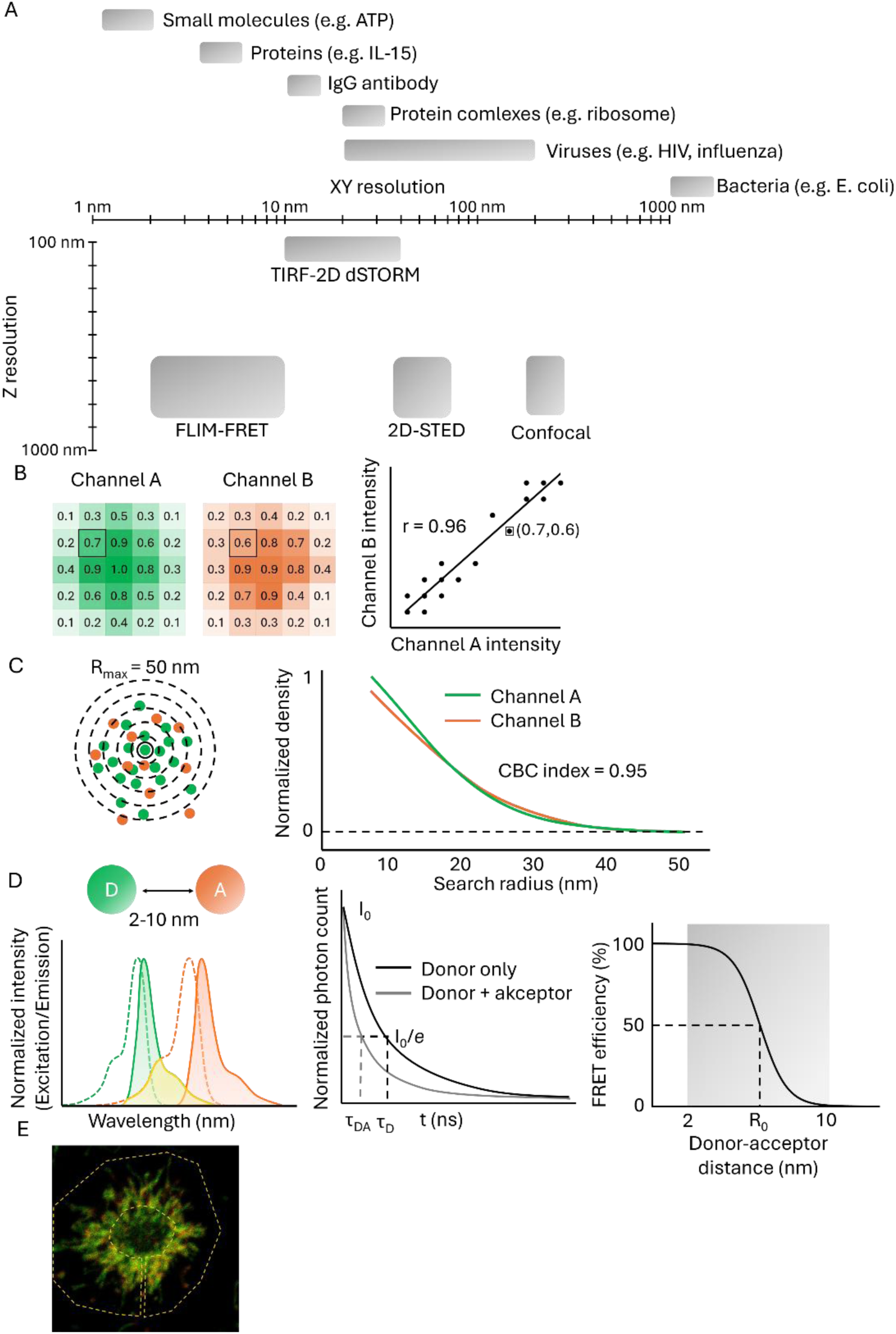
Conceptual overview of the resolution and principles of fluorescence-based proximity measurements. **(A)** Approximate sizes of selected biological structures and the lateral and axial resolution ranges of the microscopy approaches used here. **(B)** Intensity-based colocalisation analysis. Fluorescence intensity distributions from two proteins measured in channels A and B are compared on a pixel-by-pixel basis, and the strength and direction of their potential linear relation is quantified using Pearson’s correlation coefficient. **(C)** Coordinate-based colocalisation analysis. Local point densities are evaluated around detections in the two channels within a defined search radius (*R_max_*); similarity between the resulting density profiles is expressed as the CBC index. **(D)** FRET measures molecular proximity at distances of approximately 2–10 nm. Spectral overlap enables energy transfer from donor to acceptor, resulting in a reduced donor fluorescence lifetime and a distance-dependent FRET efficiency determined by the Förster radius (*R_0_*). **(E)** Representative dual-colour fluorescence image illustrating the selection of a region of interest (ROI) containing microvilli for confocal, STED and FLIM–FRET analyses.

Confocal and STED microscopy revealed significant colocalisation of IL-15Rα with both MHC class I and MHC class II molecules on B-cell microvilli. Pearson’s correlation coefficients were significantly higher for the IL-15Rα – MHC class I and IL-15Rα – MHC class II pairs than for the negative control, indicating that both MHC classes were non-randomly distributed within the same membrane regions. The stronger correlation observed for MHC class II than for MHC class I was consistent across the imaging approaches. The lower Pearson’s correlation coefficients obtained using STED than those acquired using confocal microscopy are compatible with the greater resolving power of STED, which separates membrane structures that appear merged at the diffraction-limited resolution of confocal microscopy. Thus, the confocal measurements indicate a shared membrane distribution, whereas the STED measurements provide evidence that this co-distribution persists at a finer spatial scale.

Single-molecule localisation microscopy provided an additional level of spatial information. The significantly higher CBC index for MHC class II and IL-15Rα than for the negative control supports the nanoscale co-organisation of these molecules. In contrast, the difference between MHC class I and IL-15Rα did not reach statistical significance. Importantly, the CBC index should not be interpreted as a direct measure of molecular binding. A high CBC value indicates spatial co-organisation over the characteristic length scale defined by the analysis, which is substantially greater than the Förster radius for FRET measurements. Molecules located at the periphery of one structure may yield high CBC values without being in direct molecular contact. Conversely, a low or near-zero CBC value does not necessarily exclude molecular proximity, particularly when one molecular species is relatively uniformly distributed and therefore generates a flat local density profile. [23–25]. We therefore interpret the CBC measurements as evidence of nanoscale co-organisation rather than direct molecular association.

The FRET measurements provide complementary evidence at a substantially smaller spatial scale. MHC class II exhibited higher FRET efficiency with IL-15Rα than did MHC class I, consistent with the stronger colocalisation of MHC class II and IL-15Rα observed by confocal and STED microscopy. The difference in FRET efficiency may reflect differences in molecular density and organisation, the size and composition of membrane nanodomains, or the relative accessibility and orientation of the fluorescent labels.

Our findings suggest that this organisation may have functional significance. Because microvilli are positioned to mediate the earliest membrane contacts between lymphocytes, the enrichment of IL-15Rα together with MHC class I and MHC class II may enable a B cell to present multiple classes of information to an approaching T cell within the same specialised membrane protrusions. Rather than requiring the independent recruitment of IL-15Rα and MHC molecules only after immunological synapse formation, their pre-existing spatial organisation on microvilli could facilitate the rapid establishment of productive cell–cell contacts and the subsequent assembly of the immunological synapse. This interpretation is consistent with our previous observation that IL-15Rα and MHC class II can be jointly recruited to the immunological synapse and that IL-15 trans-presentation and antigen presentation can influence one another. The present findings extend these observations both temporally and spatially by indicating that their coordinated organisation is already detectable on B-cell microvilli before synapse formation.

The association of IL-15Rα with MHC class II is particularly remarkable. Whereas MHC class I-mediated antigen presentation primarily engages CD8+ T cells, MHC class II-mediated antigen presentation engages CD4+ T cells. IL-15 trans-presentation has traditionally been considered predominantly in the context of CD8+ T-cell and NK-cell activation. [12]. The presence of IL-15Rα in close spatial proximity to MHC class II on B-cell microvilli therefore raises the possibility that IL-15 trans-presentation and MHC class II-mediated antigen presentation may be functionally integrated during B-cell–CD4+ T-cell interactions [13]. Whether this organisation contributes to CD4+ T-cell activation, differentiation or effector function remains to be established. In conclusion, our multiscale imaging data reveal a previously unrecognised spatial organisation of IL-15Rα with MHC class I and MHC class II on B-cell microvilli. The convergence of diffraction-limited and super-resolution imaging, single-molecule spatial analysis and FRET indicates that these molecules occupy shared membrane environments and, particularly in the case of IL-15Rα and MHC class II, exhibit nanoscale proximity. We propose that B-cell microvilli may function as pre-synaptic membrane platforms in which the molecular machinery required for IL-15 trans-presentation and antigen presentation is spatially organised before stable T-cell engagement. Such an organisation could facilitate the rapid integration of cytokine trans-presentation, antigen recognition and adhesion during the earliest stages of B-cell–T-cell communication. Directly testing whether disruption of this microvillar organisation alters the efficiency of IL-15 trans-presentation or antigen presentation will be an important direction for future studies.

## Materials and methods

### Cell culture

Human Raji B cells (Burkitt lymphoma cell line; RRID: CVCL_0511) were used in this study and were retrovirally transduced to express a HaloTag–IL-15Rα fusion protein, as described previously [15]. Cells were cultured in RPMI 1640 medium (Sigma-Aldrich, Merck KGaA, Darmstadt, Germany) supplemented with 10% (v/v) heat-inactivated FBS, GlutaMAX (Thermo Fisher Scientific Inc., Waltham, MA, USA) and Geneticin (Sigma-Aldrich, Merck KGaA, Darmstadt, Germany) as a selective agent. Cells were grown in a humidified atmosphere containing 5% CO₂ at 37 °C.

### Immunofluorescence labelling

#### Immunofluorescence labelling for confocal and STED microscopy

For STED imaging, clean poly-L-lysine-coated coverslips were prepared as detailed in the Supplementary Information (SI). Raji–IL-15Rα–HaloTag cells were labelled with HaloTag STAR RED ligand (abberior GmbH, Göttingen, Germany), STAR 580-conjugated anti-MHC class I Fab fragment (ST580-W6/32 Fab), STAR 580-conjugated anti-MHC class II Fab fragment (ST580-L243 Fab), STAR RED-conjugated anti-TfR antibody (STRED-MEM75 mAb) or STAR RED-conjugated anti-β2-microglobulin Fab fragment (STRED-L368 Fab). Fab fragment preparation and labelling are detailed in the Supplementary Information (SI). Cells were washed in HBSS and labelled with 0.5 μM HaloTag STAR RED ligand or 0.05 mg/ml antibody for 20 min at 37 °C in HBSS. After labelling, cells were washed twice and plated in 100 μl HBSS onto the coated coverslips and incubated in the dark for 20 min to allow adhesion. Cells were fixed by adding 50 μl of 6% formaldehyde solution, resulting in 2% final final concentration. Microscope slides were washed in 96% ethanol and allowed to dry on paper towels. Abberior Mount Solid Antifade (abberior GmbH, Göttingen, Germany) was pre-warmed to room temperature. If the mounting medium had solidified, the tube was heated in a water bath at 50 °C for 15 min. Subsequently, 15 μl of mounting medium was pipetted onto the microscope slides. Coverslips were placed with cell-side down into the mounting medium and allowed to dry overnight at 4 °C.

#### Immunofluorescence labelling for STORM

Raji-HaloTag–IL15Rα cells were washed in HBSS (1200 RPM, 5 min, room temperature), then labelled with 5 µM HaloTag FLUX 640 ligand (abberior GmbH, Göttingen, Germany) and Fab fragments or complete mAbs at 0.05 mg/ml concentration in 50 µl volume. Cells were labelled with the following antibody pairs: A647-MEM75 mAb (anti-TfR) with Cy3B-W6/32 Fab (anti-MHC class I) as a negative control; HaloTag FLUX 640 ligand with either Cy3B-W6/32 Fab or Cy3B-L243 Fab (anti-MHC class II). As a positive control, Cy3B-L243 Fab was combined with A647-W6/32 Fab. After 20 min incubation at 37 °C in HBSS, cells were washed twice with HBSS. 100 µl cells were plated on poly-L-lysine coated 8-well ibidi chambers (ibidi GmbH, Gräfelfing, Germany). After 20 minutes incubation, 50 μl 6% formaldehyde solution was added for fixation resulting in 2% final concentration.

#### Immunofluorescence labelling for FLIM-FRET measurement

Raji-HaloTag–IL15Rα cells were washed in HBSS, then labelled with 5 µM HaloTag TMR ligand (G8251, Promega Corporation, Madison, WI, USA) and 0.05 mg/ml antibodies in 50 µl volume. The following labels were applied to the cells: HaloTag TMR ligand alone or with A647-MEM75 mAb (anti-TfR); HaloTag TMR ligand alone or with A647-W6/32 mAb (anti-MHC class I); HaloTag-TMR ligand alone or with A647-L243 mAb (anti-MHC class II); A546-L368 mAb (anti-β2-microglobulin) alone or with A647-W6/32 mAb. After 20 min incubation at 37°C in HBSS, cells were washed twice. Finally, cells were suspended in 200 µl Leibovitz’s L-15 imaging medium (Thermo Fisher Scientific Inc., Waltham, MA, USA) and seeded in poly-L-lysine coated 8-well ibidi chambers (ibidi GmbH, Gräfelfing, Germany) for 15 minutes at room temperature before the measurement.

### Imaging methods

#### Confocal and STED imaging and colocalisation analysis

Imaging was carried out on a Zeiss Axiovert 200M microscope (Carl Zeiss Microscopy GmbH, Jena, Germany) with STEDYCON microscope upgrade (abberior Instruments GmbH, Göttingen, Germany) equipped with a Zeiss α Plan-Apochromat 100x/1.46 Oil DIC immersion objective (Carl Zeiss Microscopy GmbH, Jena, Germany). The samples were labelled with STAR RED and STAR580 dyes, which was excited with 640 and 561 nm lasers, in conjunction with a STED depletion laser at 775 nm. The emission was detected with 660 and 604 nm detectors. The resolution was set to 54 nm.

The colocalisation analysis was performed in the MATLAB environment using code written by Prof. Péter Nagy. The MATLAB code performs the following steps:

1. The batchPreProcess programme is opened in MATLAB, and the input folder is selected.
2. The ROI is selected manually on merged images.
3. A Gaussian filter is applied to the images, and thresholding is performed manually.
4. Pearson’s correlation coefficient is calculated between the two fluorescence channels to quantify the linear relationship between their intensity distributions.

Further details are provided in the Supplementary Information (SI).

#### Image acquisition for STORM

The imaging setup was based on an Olympus IX73 microscope body (Evident Corporation, Tokyo, Japan) equipped with a high-NA objective (Olympus UPlanApo 100×/1.5 NA, Evident Corporation, Tokyo, Japan) and an sCMOS camera (ORCA-Fusion, Hamamatsu Photonics K.K., Hamamatsu, Japan). Furthermore, it featured red and green excitation lasers (640 and 532 nm lasers; 100 mW nominal laser power; OBIS Laser Box, Coherent, Inc., Santa Clara, CA, USA). The setup was operated in spinning TIRF excitation mode (iLas 2, Gataca Systems SAS, Massy, France), yielding an excitation intensity of 1 kW/cm^2^. The lasers were filtered using a quad dichroic mirror (Semrock Di01-R405/488/532/635, IDEX Health & Science LLC, West Henrietta, NY, USA) and an emission filter (ZET405/488/532/642m, Chroma Technology Corp., Bellows Falls, VT, USA) placed in the upper deck of the microscope body. The emission was split into green and red channels using a Laser Beamsplitter H 643 LPXR Superflat, 650/SP BrightLine HC Shortpass Filter and 690/70 H Bandpass Filter (AHF analysentechnik AG, Tübingen, Germany), and was imaged side by side on the camera chip. Illumination and image acquisition were controlled by VisiView^®^ (Visitron Systems GmbH, Puchheim, Germany). To maintain constant focus during imaging, a focus-hold system was built using an IR laser diode (785 nm), which was totally internally reflected at the coverslip surface. The reflected beam was then captured by an sCMOS camera (Basler AG, Ahrensburg, Germany). Any vertical (z-axis) movement of the sample caused a lateral shift of the beam on the camera chip. A z-piezo positioned beneath the objective was used to correct for z-drift. TetraSpeck 100 nm fluorescent beads (T7279, Thermo Fisher Scientific Inc., Waltham, MA, USA) were added to the sample as fiducial markers. The beads were sonicated in an ultrasonic bath for 2 min, diluted 1:100,000 in PBS and applied to the sample for 5 min. Afterwards, the sample was rinsed with PBS. For dSTORM imaging, PBS was exchanged for blinking buffer consisting of 10% glucose, 500 μg/ml glucose oxidase, 40 μg/ml catalase and 50 mM cysteamine in PBS (pH 7.5). The dSTORM buffer was prepared freshly immediately before imaging or replaced between measurements, as its pH tends to shift after 1 h, potentially adversely affecting imaging quality. For each cell, 10,000 frames were recorded at a frame rate of 50 Hz (20 ms exposure time).

#### Evaluation for STORM data

The localisations were fitted using the ThunderSTORM plugin [26]. The localisations of the fiducial markers were then used to correct lateral drift and to register the green and red channels using the sdt-python library.

STORM images were reconstructed starting from frame 500. A region of interest (ROI) was applied to the centrally located, homogeneously illuminated region of the reconstructed images. Duplicate localisations were then removed; that is, localisations closer to each other than their localisation uncertainty were merged.

A density filter was subsequently applied to remove all localisations with fewer than three neighbours within a 50 nm radius. The coordinate-based colocalisation index was calculated on the resulting images.

Colocalisation between the two channels was quantified at the single-molecule level using coordinate-based colocalisation (CBC) analysis, which operates on localisation coordinates rather than reconstructed pixel images and therefore avoids the intensity-based assumptions of Pearson’s and Manders’ coefficients. For each localisation *A_i_* of the base channel *A*, the number of neighbouring localisations in its own channel and in the second channel *B* was counted within concentric circles of increasing radius *r*, sampled at *j* equally spaced values up to *R_max_ = 50 nm*. The counts for the base channel *A* were normalised to the corresponding density at *R_max_*, according to

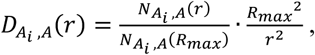

where *D_Ai,A_(r)* is the distribution of localisations from channel *A* around localisation *A_i_*; *N_Ai,A_(r)* is the number of localisations in channel *A* within radius *r* around localisation *A_i_*; and *N_Ai,A_(R_max_)* is the total number of localisations in channel *A* within radius *R_max_* around localisation *A_i_*. The distribution of localisations from the second channel was calculated using an analogous expression:

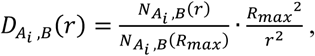

where *D_Ai,B_(r)* is the distribution of localisations from channel *B* around localisation *A_i_*; *N_Ai,B_(r)* is the number of localisations in channel *B* within radius *r* around localisation *A_i_*; and *N_Ai,B_(R_max_)* is the total number of localisations in channel *B* within radius *R_max_* around localisation *A_i_*. For a spatially uniform distribution, the density profile is expected to remain constant at *D(r) = 1* across all radii *r*. The similarity of the two resulting density profiles was evaluated using the Spearman’s rank correlation coefficient S_Ai_, which is insensitive to differences in absolute molecular density between channels because it compares the rank order, rather than absolute magnitude. Spearman’s rank correlation coefficient was calculated using the following equation:

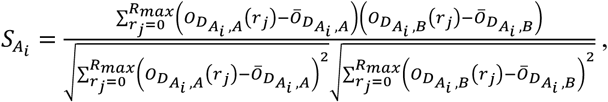

where *O_DAi,A_(r_j_)* and *O_DAi,B_(r_j_)* are the rank-transformed values of the *D_Ai,A_(r)* and *D_Ai,B_(r)*, respectively, at the sampled radius *r_j_*; and O̅_DAi,A_ and O̅_DAi,B_ are the mean ranks of the corresponding density profiles across all sampled radii. Thus, *S_Ai_* is the Pearson’s correlation coefficient calculated between the ranked density profiles, i.e., the Spearman’s rank correlation coefficient. The coefficient was subsequently weighted by the distance *E_(Ai,B)_* from *A_i_* to its nearest neighbour in the other channel, yielding the colocalisation value:

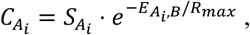

where *C_Ai_* is the coordinate-based colocalisation (CBC) value assigned to localisation *A_i_*; *S_Ai_* is the Spearman’s rank correlation coefficient between the density profiles of channel *A* and *B* around localisation *A_i_*; *e* is the Euler’s number; and *E_Ai,B_* is the Euclidean distance from *A_i_* to its nearest neighbour in the channel *B*. Localisations without a cross-channel neighbour within *R_max_* were assigned *C_Ai_ = 0*. The resulting index ranges from −1 for mutually exclusive (segregated) distributions, through 0 for spatially independent distributions, to +1 for perfectly correlated distributions. The analysis was performed reciprocally, with each channel serving as the base channel, and the two values were averaged. [23]

#### FLIM-FRET measurement and evaluation

To measure interactions between proteins on B-cell microvilli, FLIM-FRET was used. The measurements were carried out on a Nikon A1 confocal microscope (Nikon Corporation, Tokyo, Japan) equipped with a Plan-Apochromat 60× water-immersion objective (NA = 1.27; Nikon Corporation, Tokyo, Japan) and a time-correlated single-photon counting (TCSPC) extension (PicoQuant GmbH, Berlin, Germany). The pinhole size was 1.2 Airy units, while the zoom factor was set to 4.5. Before the measurements, preview confocal images were recorded using 561 nm and 647 nm continuous-wave (CW) lasers and detected through 570/620 nm and 663/738 nm bandpass filters to visualise A546- and A647-labelled proteins, respectively. For all FLIM measurements, the zoom factor was kept constant. Therefore, cropped confocal images and FRET maps are shown in Fig. 3 to improve visualisation, while the corresponding uncropped images are provided in Fig. S2.

To excite the HaloTag TMR ligand or A546 during FLIM measurements, a 565 nm picosecond pulsed laser with a repetition rate of 20 MHz was used. Emission was detected through a 600/50 nm emission filter using a PMA Hybrid 40 photon-counting photomultiplier (PicoQuant GmbH, Berlin, Germany). Data were collected for 60 s for the HaloTag TMR or A546 donor. All measurements were performed at room temperature (23 °C).

The fluorescence decay curves were analysed using SymphoTime 64 software (PicoQuant GmbH, Berlin, Germany). First, the entire image was intensity-thresholded to exclude background pixels. Next, a freehand-drawn ROI was applied to pixels containing the microvilli of a single cell. The fluorescence intensity decay curves of the selected areas were fitted with two lifetime components using a multiexponential reconvolution model:

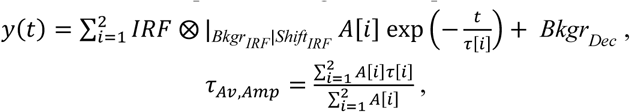

where *τ[i]* and *A[i]* are the exponential decay time and amplitude of the i^th^ component, respectively; *Bkgr_IRF_* is the background correction for the instrument response function (IRF); *Shift_IRF_* is the correction for temporal IRF displacement; *Bkgr_Dec_* is the background correction; and *τ_Av,Amp_* is the amplitude-weighted average lifetime.

To calculate the FRET efficiency (*E*) within the defined ROI, the following equation was used:

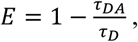

where *τ*_DA_ and *τ*_D_ are the amplitude-weighted average fluorescence lifetimes of the donor in the presence and absence of the acceptor, respectively.

For selected cells, the pixel-by-pixel distribution of FRET efficiency and the corresponding FRET-efficiency histogram were also calculated within the defined ROI.

#### Statistics

Outliers were identified using the robust regression and outlier removal (ROUT) method implemented in GraphPad Prism 10, with the false discovery rate (Q) set to 1%. Replicate values within each row were averaged before outlier detection, and data points flagged as outliers by the ROUT procedure were excluded from further analysis.

To assess distributional assumptions, the normality of residuals was evaluated using multiple tests implemented in GraphPad Prism 10, including the Anderson–Darling, D’Agostino–Pearson omnibus, Shapiro– Wilk and Kolmogorov–Smirnov tests (α = 0.05).

For comparisons of more than two groups, data were analysed using Brown–Forsythe and Welch’s heteroscedastic one-way ANOVA (GraphPad Prism 10), which does not assume equal variances across groups. When the omnibus test was significant, pairwise differences between group means were assessed using Welch’s t-tests, and P values were adjusted for multiple comparisons using the two-stage linear step-up procedure of Benjamini, Krieger and Yekutieli to control the false discovery rate (Q = 0.10).

If the data did not meet the assumption of normality, group medians were compared using the Kruskal– Wallis test (GraphPad Prism 10). Significant omnibus results were followed by Dunn’s post hoc test, comparing the mean rank of each group with that of the control; P values were adjusted for multiple comparisons using a family-wise significance threshold of 0.001.

## Supplementary Information

The online version contains supplementary information.

## Acknowledgments

We thank Edina Nagy for technical assistance and Prof. Péter Nagy for providing the MATLAB routine for the evaluation of STED images. Microscopy was carried out at the Sándor Damjanovich Cell Analysis Core Facility of the University of Debrecen (Cellular BioImaging Hungary, Euro-BioImaging Node). This work was supported by the following grants: ANN 135107, K146028, 2024-1.2.2-ERA_NET-2024-00009 from the National Research, Development and Innovation Office, Hungary and the University of Debrecen Program for Scientific Publication. The study was supported by the Austrian Science Fund (FWF) (10.55776/I5056). For open access purposes, the author has applied a CC BY public copyright license to any author-accepted manuscript version arising from this submission. This research work was conducted with the support of the National Academy of Scientist Education Program of the National Biomedical Foundation.

## Author contributions

**István Rebenku**: Conceptualization, Methodology, Formal analysis, Investigation, Writing - Original Draft, Writing - Review & Editing, Visualization. **Noémi Bilakovics**: Methodology, Formal analysis, Investigation, Writing - Original Draft, Writing - Review & Editing, Visualization. **Levente Szolyka**: Methodology, Formal analysis, Investigation, Writing - Original Draft, Writing - Review & Editing, Visualization. **Lukas Velas**: Methodology, Formal analysis, Investigation. **Tamás Garda**: Methodology. **Gerhard J. Schütz**: Methodology, Resources, Review & Editing, Funding acquisition. **György Vámosi**: Conceptualization, Methodology, Writing - Original Draft, Writing - Review & Editing, Visualization, Supervision, Project administration, Funding acquisition.

## Data availability

This study includes no data deposited in external repositories. All data supporting the findings of this study are available within the paper and its Supplementary Information.

## Supplementary figures and legends

**Fig. S1.**
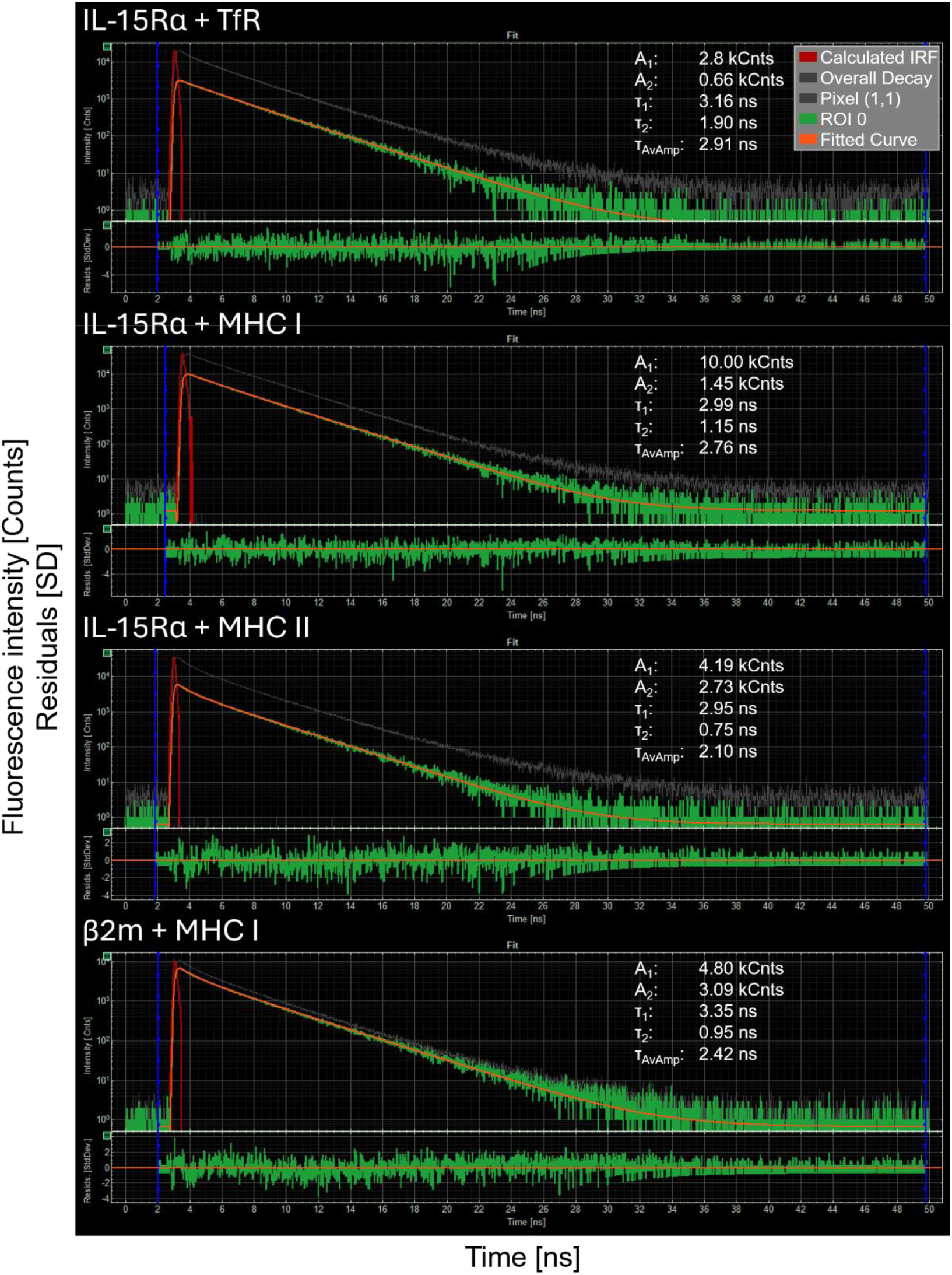
Fluorescence intensity decay curves from FLIM-FRET measurements on Raji-cell microvilli. The panel shows the fluorescence intensity lifetime decay curves of the TMR (1^st^-3^rd^ rows) and A546 (4^th^ row) donors. 1^st^ row: TMR-tagged HaloTag-IL-15Rα and A647-labelled transferrin receptor (TfR); 2^nd^ row: TMR-tagged HaloTag-IL-15Rα and A647-labelled MHC class I; 3^rd^ row: TMR-tagged HaloTag-IL-15Rα and A647-labelled MHC class II; 4^th^ row: A546-labelled MHC class I and A647-labelled β2-microglobulin (β2m). The raw data (green) were fitted with two lifetime decay components (orange), characterized by the preexponential amplitudes A1 and A2 and the fluorescence lifetimes τ1 and τ2. The amplitude-weighted average lifetime of the donor, τAv,Amp, was used to calculate the FRET efficiency. Each decay curve represents photon counts from the hand-drawn ROI of a selected cell. The fit residuals are shown below the decay curves. The instrument response function is represented by the peaked curve at the beginning of the decay (red), calculated using the SymphoTime 64 software.

**Fig. S2.**
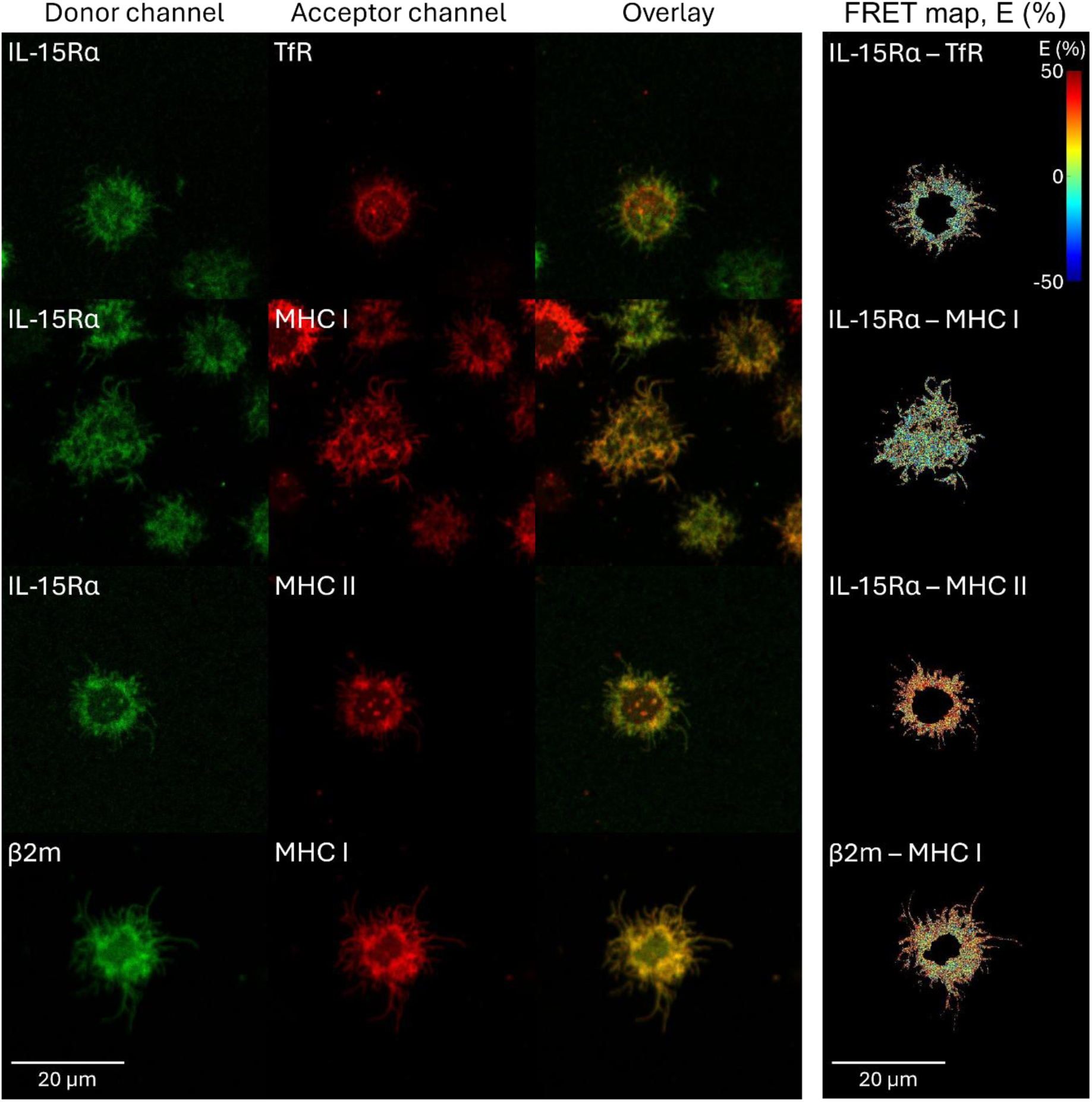
Uncropped confocal images and FRET maps from FLIM–FRET measurements of Raji-HaloTag– IL-15Rα cells. Representative uncropped confocal images corresponding to the cropped images presented in Fig. 3 are shown. Columns show the FRET donor channel, acceptor channel, merged image, and FRET-efficiency map, respectively. The first row shows a cell labelled with HaloTag TMR ligand and A647-MEM75 (anti-transferrin receptor) mAb; second row: HaloTag TMR ligand and A647-W6/32 (anti-MHC class I); third row: HaloTag TMR ligand and A647-L243 (anti-MHC class II); fourth row: A546-W6/32 (anti-MHC class I) and A647-L368 (anti- β2-microglobulin) mAbs. The images in Fig. 3A were cropped from these original fields of view to facilitate visualization of the microvillar regions. Scale bar: 20 µm.

## Supplementary methods

### Coverslip coating

Round glass coverslips (12 mm diameter) were cleaned sequentially in 1% Hellmanex™ III solution (Hellma GmbH & Co. KG, Müllheim, Germany) and 100 mM HCl for 60 min each. Coverslips were rinsed thoroughly with ultrapure water (Milli-Q^®^, Merck KGaA, Darmstadt, Germany) after each cleaning step and air-dried on lens-cleaning tissue (Grade 105, Whatman™, Cytiva, Global Life Sciences Solutions USA LLC, Marlborough, MA, USA) protected from dust. For coating, each coverslip was incubated with 100 µl of 1 mg/ml poly-L-lysine solution for 30 min. Excess solution was removed by gently touching the edge of each coverslip with lens-cleaning tissue. Coverslips were rinsed with ultrapure water and air-dried on lens-cleaning tissue protected from dust. Poly-L-lysine-coated 8-well ibidi chambers (ibidi GmbH, Gräfelfing, Germany) were prepared by incubating the wells with 200 µl of 1 mg/ml poly-L-lysine for 30 min at room temperature, washing them twice with ultrapure water, and allowing them to air-dry while protected from dust.

### Fab fragment preparation and antibody labelling

The W6/32 (RRID: CVCL_7872), L243 (RRID: CVCL_4533), and L368 (RRID: CVCL_E981) monoclonal antibodies were prepared from hybridoma cell-culture supernatants. The MEM75 antibody (cat. no. 11-235-C100) was purchased from EXBIO Praha, a.s. (Vestec, Czech Republic). For antibody purification, hybridoma cell-culture supernatants were clarified by centrifugation at 1,230 × g for 20 min at 4 °C and stored at −20 °C until purification. After thawing and cooling to 4 °C, antibodies were purified by Protein A affinity chromatography using a Protein A–Sepharose™ 4 Fast Flow column (P9424, Sigma-Aldrich, Merck KGaA, Darmstadt, Germany) connected upstream of a Sepharose™ CL-4B guard column (CL4B200, Sigma-Aldrich, Merck KGaA, Darmstadt, Germany). The columns were equilibrated and washed with phosphate buffer (pH 8.0). Bound antibodies were eluted from the Protein A column with citrate buffer (pH 3.7) and immediately neutralised with 1 M Tris-HCl (pH 8.5). Protein-containing fractions were pooled, diluted with PBS, and concentrated using 30-kDa molecular-weight cut-off (MWCO) Amicon® Ultra-15 centrifugal filter units (UFC9030, Millipore, Merck, Germany). The samples were washed twice with PBS, and the final antibody concentration was determined by absorbance at 280 nm using a NanoDrop 1000 spectrophotometer (Thermo Fisher Scientific Inc., Waltham, MA, USA). Sodium azide was added to a final concentration of 0.02% (w/v), and the purified antibodies were stored at 4 °C.

Fab fragments were prepared from purified monoclonal antibodies using the Pierce™ Fab Preparation Kit (cat. no. 44985, Thermo Fisher Scientific Inc., Waltham, MA, USA) according to the manufacturer’s instructions. Briefly, antibodies were buffer-exchanged into Fab Digestion Buffer, digested with immobilised papain at 37 °C, and purified using NAb™ Protein A Plus Spin Columns. Fab-containing flow-through and wash fractions were pooled and buffer-exchanged into PBS using 10-kDa molecular-weight cut-off (MWCO) Amicon® Ultra-15 centrifugal filter units (UFC9010, Millipore, Merck, Germany)

Whole antibodies and Fab fragments were fluorescently labelled with NHS ester dyes. STAR 580 or STAR RED NHS esters (abberior GmbH, Göttingen, Germany) were used for STED microscopy, Alexa Fluor™ 546 (A20002, Thermo Fisher Scientific Inc., Waltham, MA, USA) or Alexa Fluor™ 647 (A37573, Thermo Fisher Scientific Inc., Waltham, MA, USA) NHS esters for FLIM–FRET measurements, and Cyanine 3B (GEPA63101, Sigma-Aldrich, Merck KGaA, Darmstadt, Germany) or Alexa Fluor™ 647 (A37573, Thermo Fisher Scientific Inc., Waltham, MA, USA) NHS esters for STORM imaging. Antibodies and Fab fragments were adjusted to approximately 2 mg/ml in PBS, and 1 M sodium bicarbonate was added to a final concentration of 100 mM. NHS ester dyes were dissolved in anhydrous DMSO immediately before use and added to the protein solutions. Labelling reactions were incubated for 50 min at room temperature in the dark with gentle rotation. Unconjugated dye was removed by size-exclusion chromatography using Sephadex^®^ G-50 (G5080, Sigma-Aldrich, Merck KGaA, Darmstadt, Germany) columns equilibrated with PBS. Protein concentration, dye concentration, and the dye-to-protein molar ratio were determined using a NanoDrop 1000 spectrophotometer (Thermo Fisher Scientific Inc., Waltham, MA, USA). Before use, all antibodies and Fab fragments were centrifuged at 31,510 × g for 20 min at 4 °C to remove potential aggregates.

### MatLab code for STED image analysis

The batchPreProcess program written by Prof. Péter Nagy uses the following MathLab extensions:

1. Bio-Formats is a software developed by Open Microscopy Environment consortium (University of Dundee and Glencoe Software Inc.) to read and write image data.
2. DIPimage toolbox is written for quantitative image analysis at Delft University of Technology.
3. Openpicture made by Prof. Péter Nagy is a simple application for opening images with Bio-Formats.

The following MatLab code had used:

*starred=squeeze(a(:,:,4));*

*star580=squeeze(a(:,:,5)); mask=imagegate(joinchannels(’rgb’,stretch(starred,5,95),stretch(star580,5,95)),’noupdate’); mask2=imageThresholding({gaussf(starred,1),gaussf(star580,1)},’manual’,’outputtype’,’image’,’overlap’,’ different’,’foreground’,{mask,mask});*

*mask2=mask2{3}==3; coloc=calcColocalization(starred,star580,100,0.95,mask2,mask2,’pearson’,false,false); exportvariable coloc*

*exportvariable mask2*

