## Supplementary Information for "MHC Molecules on B Cell Microvilli Are Spatially Associated with IL-15Rα"

The Supplementary Information contains two figures and supplementary methods.

### Supplementary figures and legends

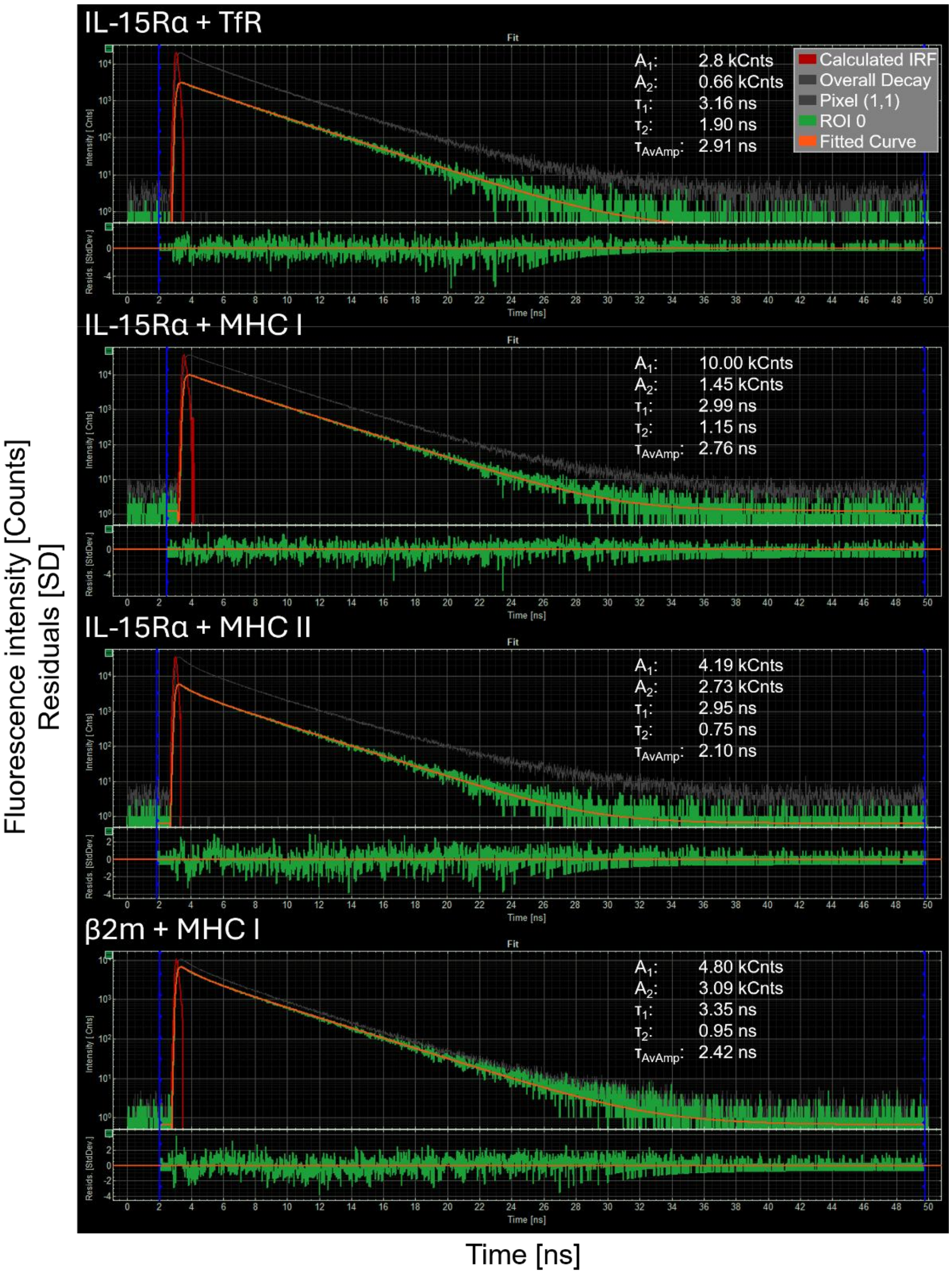


**Fig. S1. Fluorescence intensity decay curves from FLIM-FRET measurements on Raji-cell microvilli.** The panel shows the fluorescence intensity lifetime decay curves of the TMR (1st-3rd rows) and A546 (4th row) donors. 1st row: TMR-tagged HaloTag-IL-15Rα and A647-labelled transferrin receptor (TfR); 2nd row: TMR-tagged HaloTag-IL-15Rα and A647-labelled MHC class I; 3rd row: TMR-tagged HaloTag-IL-15Rα and A647-labelled MHC class II; 4th row: A546-labelled MHC class I and A647-labelled β2-microglobulin (β2m). The raw data (green) were fitted with two lifetime decay components (orange), characterized by the preexponential amplitudes $A_1$ and $A_2$ and the fluorescence lifetimes $\tau_1$ and $\tau_2$. The amplitude-weighted average lifetime of the donor, $\tau_{Av,Amp}$, was used to calculate the FRET efficiency. Each decay curve represents photon counts from the hand-drawn ROI of a selected cell. The fit residuals are shown below the decay curves. The instrument response function is represented by the peaked curve at the beginning of the decay (red), calculated using the SymphoTime 64 software.

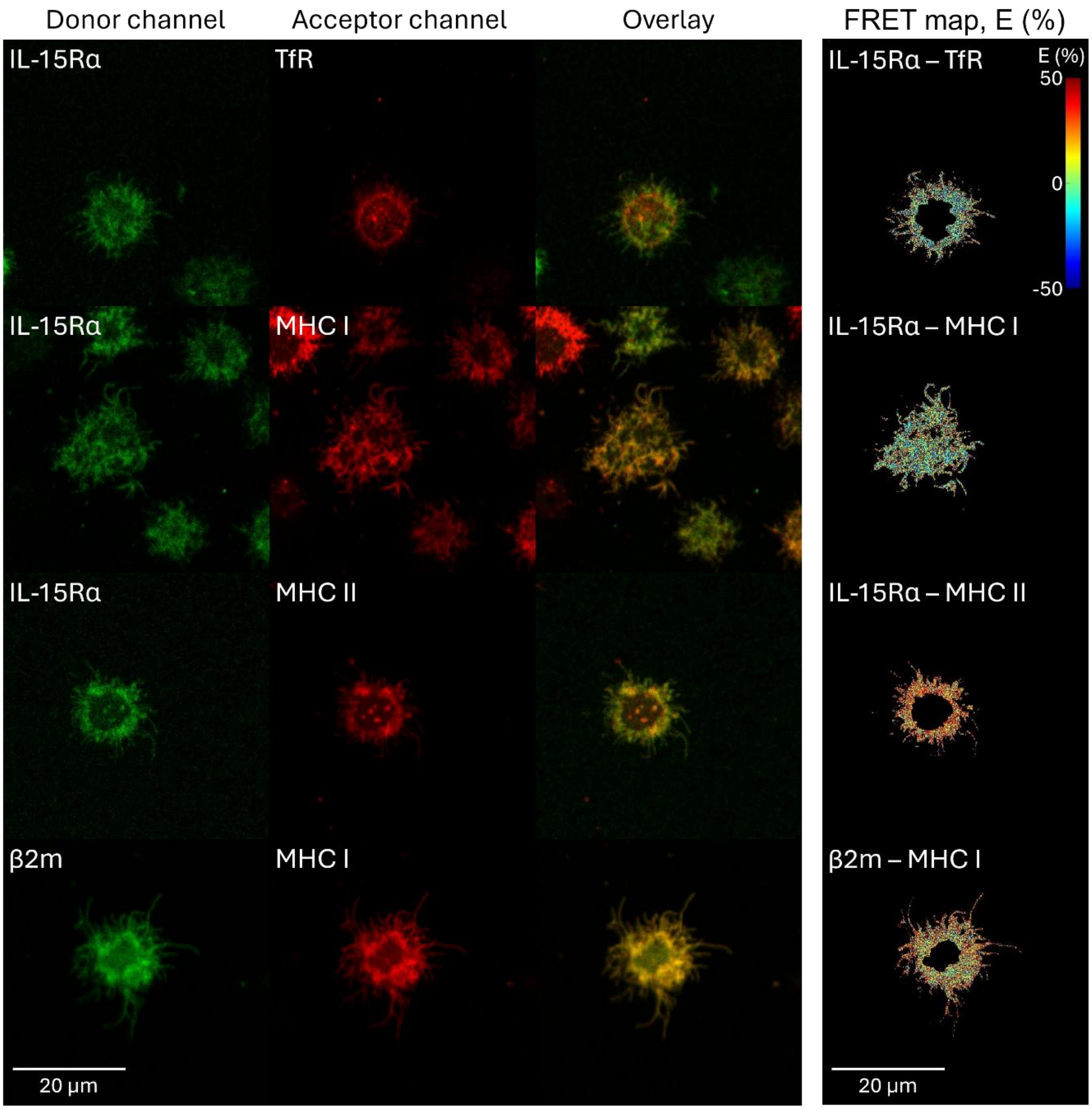


**Fig. S2. Uncropped confocal images and FRET maps from FLIM–FRET measurements of Raji-HaloTag–IL-15Rα cells.** Representative uncropped confocal images corresponding to the cropped images presented in Fig. 3 are shown. Columns show the FRET donor channel, acceptor channel, merged image, and FRET-efficiency map, respectively. The first row shows a cell labelled with HaloTag TMR ligand and A647-MEM75 (anti-transferrin receptor) mAb; second row: HaloTag TMR ligand and A647-W6/32 (anti-MHC class I); third row: HaloTag TMR ligand and A647-L243 (anti-MHC class II); fourth row: A546-W6/32 (anti-MHC class I) and A647-L368 (anti-β2-microglobulin) mAbs. The images in Fig. 3A were cropped from these original fields of view to facilitate visualization of the microvillar regions. Scale bar: 20 µm.
