## Supplementary figures and images for "MHC Molecules on B Cell Microvilli Are Spatially Associated with IL-15Rα"

### Supplementary Figure 1

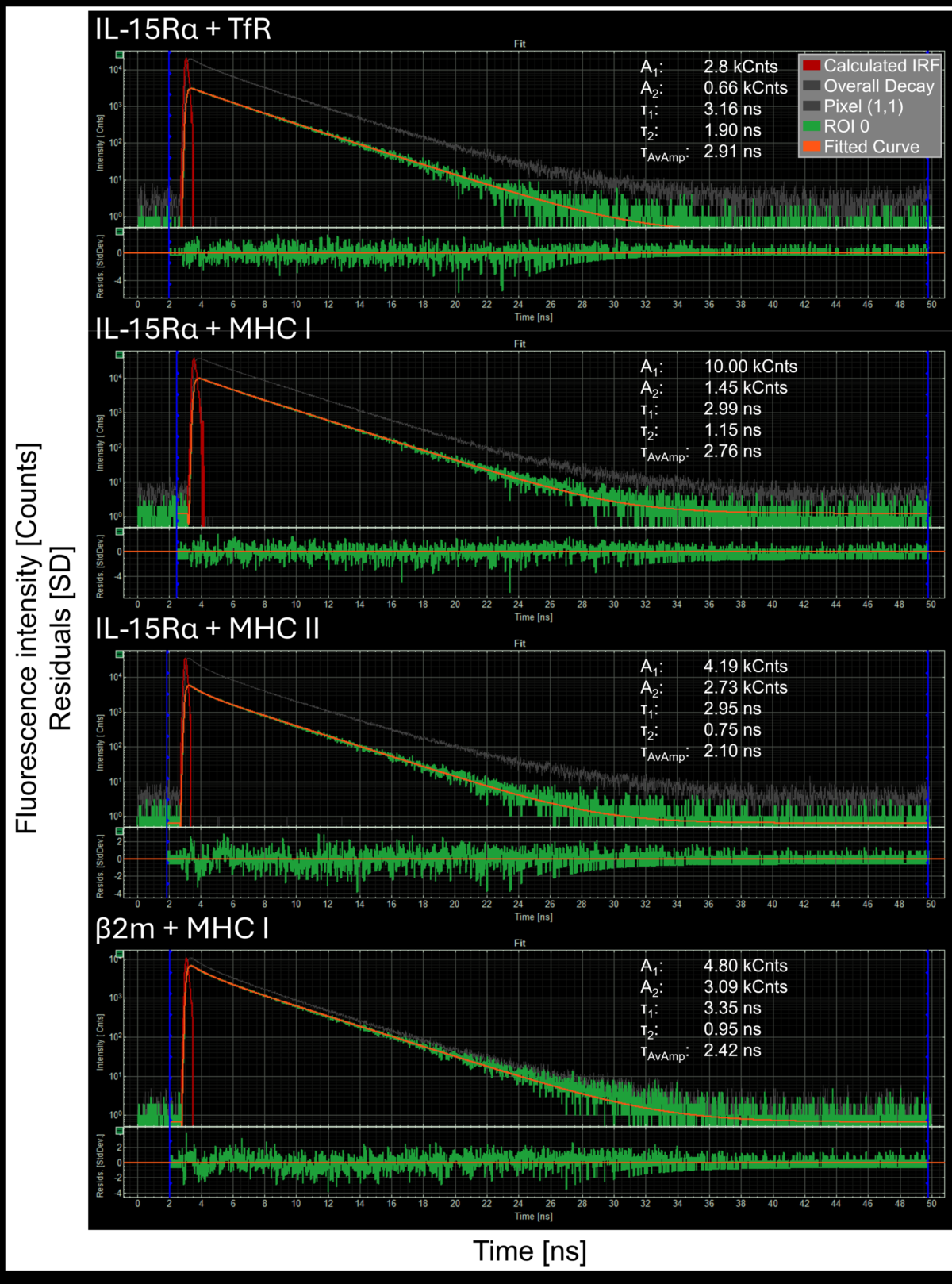

### Supplementary Figure 2

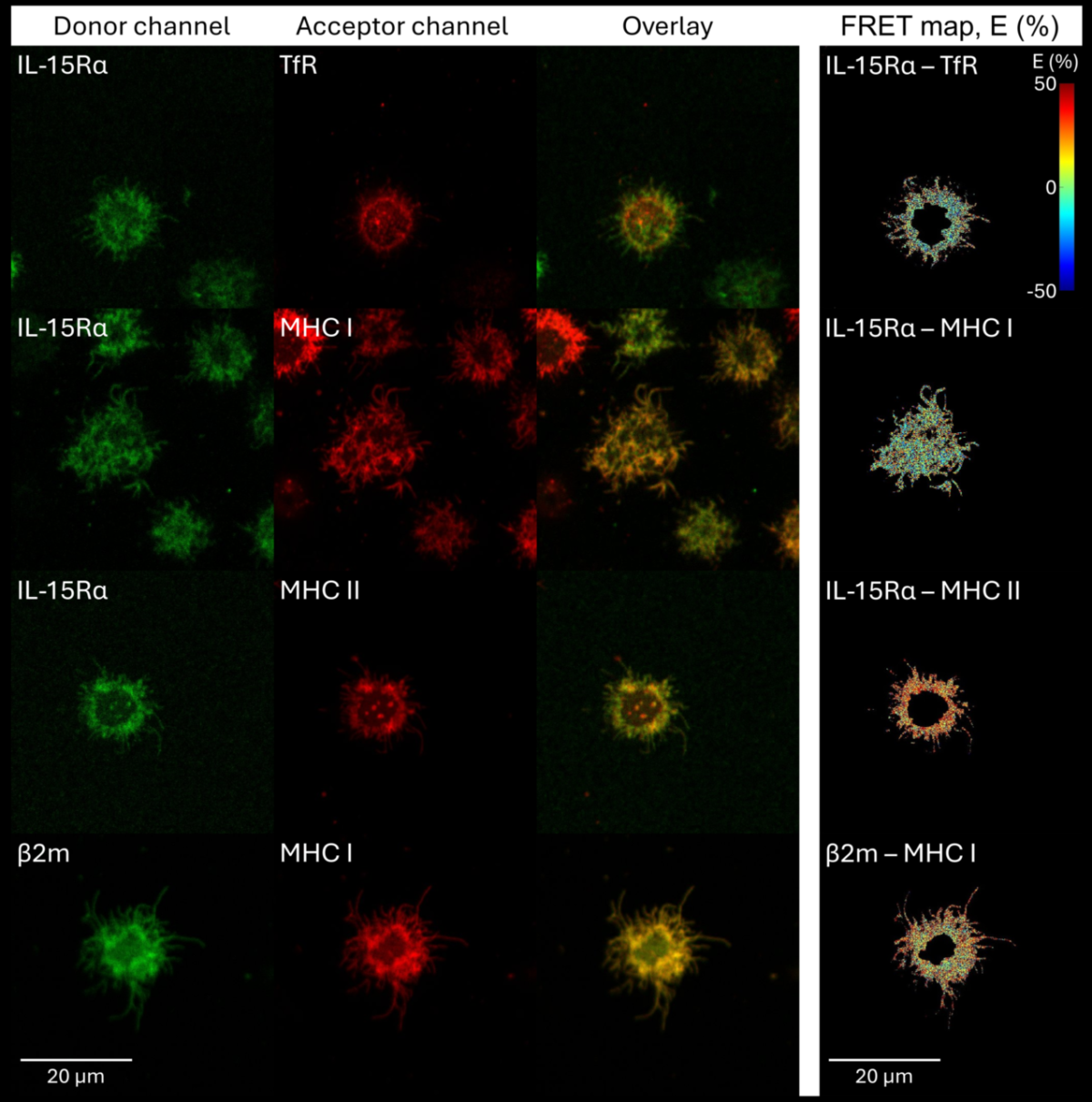
